# Characteristics of Vitelline Warbler songs and calls

**DOI:** 10.1101/2024.10.12.617988

**Authors:** Wyatt J. Cummings, David D. L. Goodman, Craig D. Layne, Katherine I. Singer, M. Whitney Thomas

**Affiliations:** Department of Biological Sciences, Dartmouth College, Hanover, NH, USA

**Keywords:** Avian vocalizations, Vitelline Warbler (*S. vitellina)*, Cayman Islands, song typology, song evolution, selection pressures, migratory species, endemic species

## Abstract

The Vitelline Warbler (*Setophaga vitellina*) is an understudied species endemic to a few small islands in the western Caribbean. Little is known beyond its phylogenetic relationship to other New World warblers. We used island-wide surveys and bioacoustic recordings to investigate the distribution, vocalizations, and ecology of *S. vitellina* across a significant portion of the species’ range on Little Cayman Island. We recorded 417 songs from 91 individuals and analyzed the length, frequency, and shape of various song components. We observed and characterized high variation in the composition and character of songs. We also describe the call of the species, document an association with gumbo limbo (*Bursera simaruba*) trees, and report interactions with other bird species. Improved knowledge of Vitelline Warblers has value for evaluating conservation threats to an island endemic and for understanding the evolution of vocalization behavior in Neotropical songbirds.

## Introduction

Birdsong is a cornerstone of ornithology, with song learning and variation providing key insights into the broader landscape of avian ecology, life history, and behavior (1–4). In North America, studies of the family Parulidae (wood-warblers) and particularly the genus *Setophaga* have yielded insights about the identification and categorization of songs (5), the effects of abiotic factors on singing behavior (6), geographical gradients of song variation (7), and advances in passive bioacoustic monitoring (8).

The Vitelline Warbler (*Setophaga vitellina*), a close relative of the Prairie Warbler (*S. discolor;* Vieillot 1809; (9)), is a poorly studied member of the *Setophaga* genus that could provide insights into Paruline evolution, speciation, and vocalizations. *S. vitellina* is a range-restricted species, found only in the Cayman Islands and the Swan Islands of the southwestern Caribbean. The warbler’s small range, currently estimated at less than 135 km^2^, has resulted in its global classification as “Near Threatened,” but little else is known about threats facing the species (10).

Little research has focused on *S. vitellina* beyond its phylogenetics and morphology. Markland and Lovette (11) conducted a thorough phylogenetic investigation of the species, confirming its recent divergence from a common ancestor *S. discolor* and establishing genetic differences between the three subspecies. Of these three subspecies, *S. vitellina vitellina* Cory 1886 is found on Grand Cayman, *S. vitellina crawfordi* Nicoll 1904 occurs on both Little Cayman and Cayman Brac, and *S. vitellina nelsoni* Bangs 1919 inhabits the Swan Islands (12). Morphology is well documented from individuals collected throughout the past two centuries. *S. vitellina* has the longest bill of any Paruline species (13). Like many *Setophaga* species, *S. vitellina* exhibits sexual dimorphism (Fig 1).

**Fig 1.**
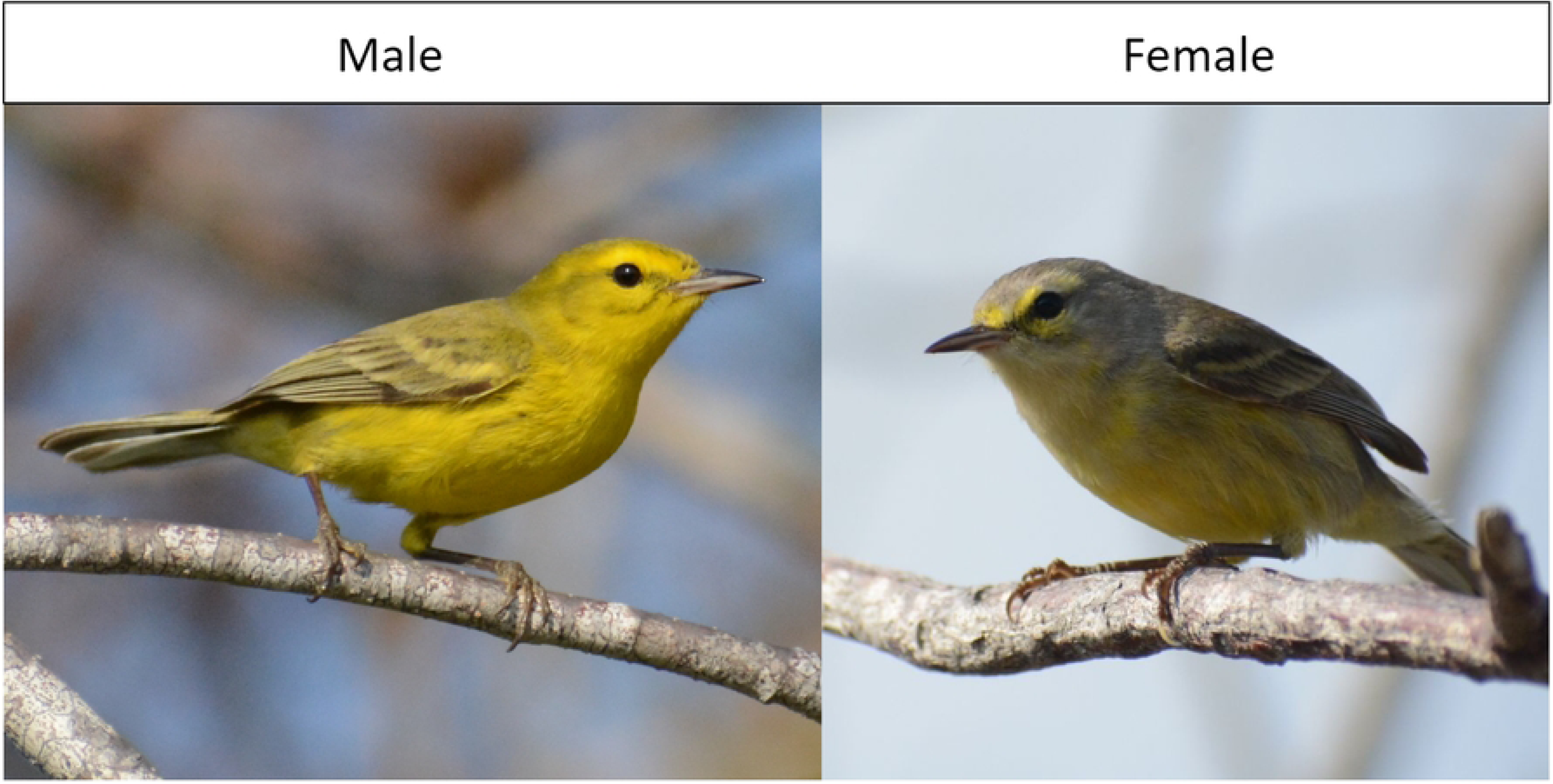
Photographs of each sex of *S. vitellina*. The male is mostly yellow with darker olive and gray-brown streaks, while the female has a mostly gray head and back, a yellow underside, and olive wings. Photos were taken on Little Cayman on March 6th, 2023 by Wyatt Cummings.

There has been no published research on the singing behavior of *S. vitellina*. Previous to this study, Cornell’s Macaulay Library had only eight audio recordings of *S. vitellina* songs (14). The song has been qualitatively compared to that of Black-throated Blue Warblers (*S. caerulescens* “Gmelin, JF” 1789), which regularly overwinters on Little Cayman (12), but no bioacoustic studies of *S. vitellina* are available to compare its vocalizations to other *Setophaga* species. Here we describe the distribution and typology of *S. vitellina* calls and songs on Little Cayman, with the aim of illuminating poorly understood aspects of the endemic songbird’s behavior.

## Materials and Methods

### Study site

We conducted field research on Little Cayman, Cayman Islands, a low-lying coral island in the western Caribbean. Little Cayman is approximately 16 km long and 1.6 km wide, making it the smallest of the three Cayman Islands. Little Cayman is also the least developed of the islands, with only 161 permanent residents as of 2021 (15). The vegetation of Little Cayman is dominated by dry forest, dry scrub, sea grape, and mangroves (16).

Little Cayman was a favorable study site for *S. vitellina* because of its low human population density and the proximity of roads to intact forest habitat. The island is circumscribed by a 35.3 km road, with three short connector roads crossing the island, for a total of 40 km of road (Fig 2).

**Fig 2.**
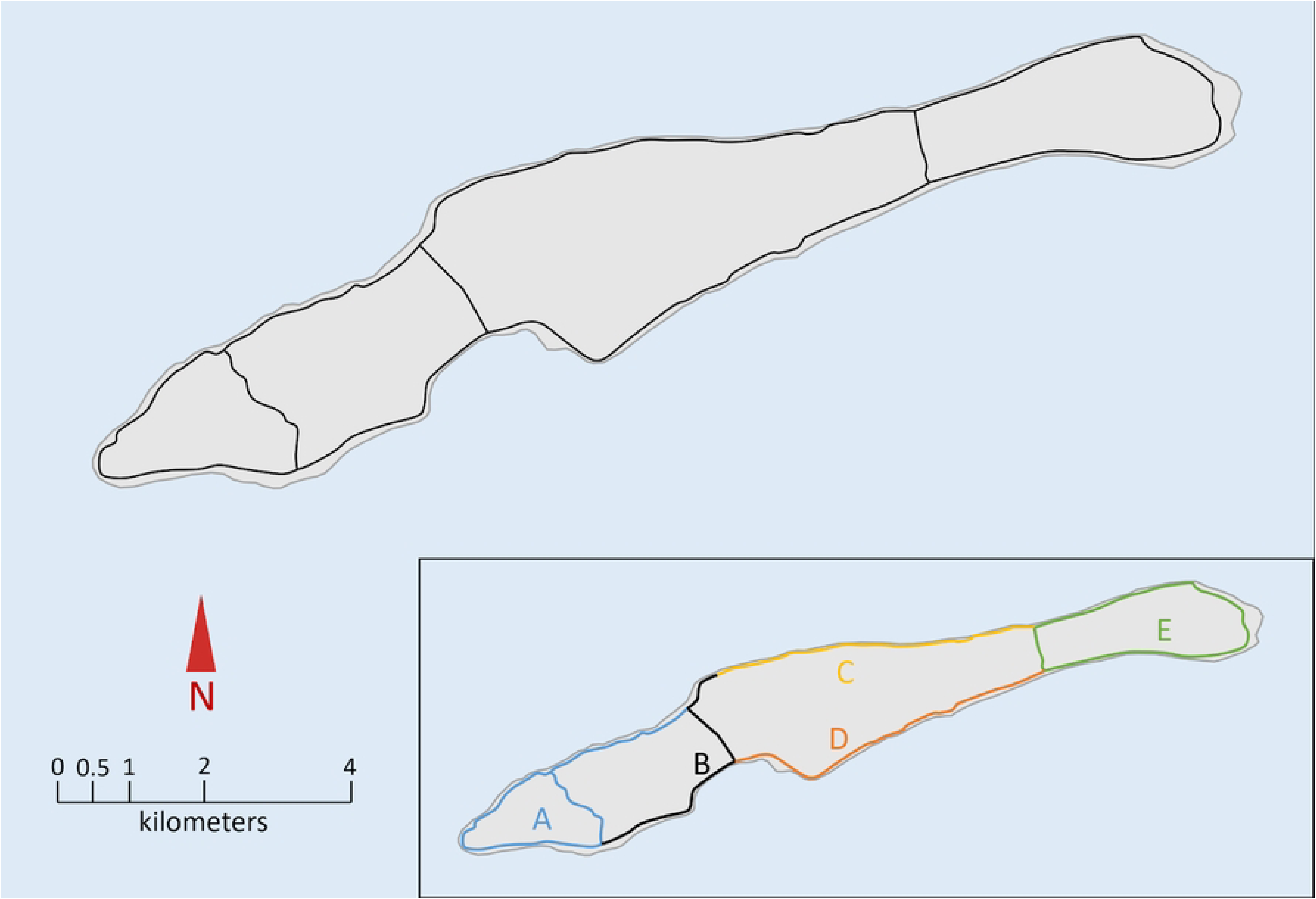
Map of the road system of Little Cayman Island. Inset demarcates five distribution survey transects routes, labeled A-E.

### Distributional surveys

We conducted distributional surveys of the entire island between 27 February and 5 March, 2023. We divided the island’s road system into five survey routes of relatively similar length (Fig 2 inset: routes A-E differentiated by color). On each survey, one or more researchers biked the entirety of the route, marking the GPS coordinates of any Vitelline Warbler heard singing using the Cornell Lab of Ornithology’s eBird GPS system, which utilizes the network assisted GPS system on the Apple iPhone. We surveyed each route once to standardize relative abundance among routes.

### Song recordings

We conducted a separate round of recording surveys from 28 February to 8 March, 2023 to collect acoustic data on as many individual warblers as possible. Each singing bird was recorded for three minutes using Merlin, a smartphone-based identification and recording app (version 2.1.5, Cornell Lab of Ornithology). We collected song recordings across the entire breadth of the island in order to account for any geographic differences in singing. We sought to avoid resampling the same individual by never repeating the same transect, but movement rates of *S. vitellina* over a 24-hour period are unknown, so we cannot rule out the possibility of duplication.

### Data analysis

All analyses of songs and calls were performed with Raven Lite (version 2.0.4). We visually categorized songs by the number of components they contained (classified as chips, up-notes, and down-notes). For each song, the time and frequency of each component was noted. Time and frequency were measured at the center of the chip’s region on the spectrogram, and at the start of the note and the end of the note for up notes and down notes. There were two note varieties (one up-note and one-down note) that we analyzed differently due to their unique shape (Fig 4). The “check” up-note declined in frequency before rising, so we measured time and frequency at the minimum frequency, as well as the start and the end of the up-note. The “parabolic” down-note rose in frequency before declining, so we measured time and frequency at the maximum frequency, as well as the start and the end of the entire down-note.

JMP Pro 16 was used for analyses of time, frequency, shape, and song structure. For each song type, and for all data combined, the Distribution feature was used to assess normality and determine the mean and standard error for each parameter. For comparison of parameter values between song types and shape varieties, we used a one-way ANOVA test.

## Results

### Distribution on Little Cayman

During our surveys, we encountered 56 singing *S. vitellina*, equivalent to 1.4 singing individuals per km of transect. Distribution surveys yielded clear qualitative data on *S. vitellina* habitat preferences. Along the island’s coast, *S. vitellina* was found in the highest densities in dry forest and scrub forest habitats (1.71 singing individuals per kilometer of transect). The species occurred in low densities in sea grape (*Coccoloba uvifera*)-dominated plant communities (0.73 singing individuals per kilometer of transect), and was almost entirely absent in areas dominated by mangroves (0.29 singing individuals per kilometer of transect).

Along the inland cross-roads, we detected the majority of observed warblers in dry forest or dry scrub (4.89 singing individuals per kilometer of transect), especially in areas with a high density of gumbo limbo (*Bursera simaruba*) trees. *Bursera simaruba* were generally infrequent along the coast, and a characteristic of elevated, dry areas on Little Cayman.

### Song characteristics

The song of *S. vitellina* is composed of three basic components (Fig 3): introductory short notes (“chips”), a long note that rises in frequency (“up-note”), and a long note that decreases in frequency (“down-note”).

**Fig 3.**
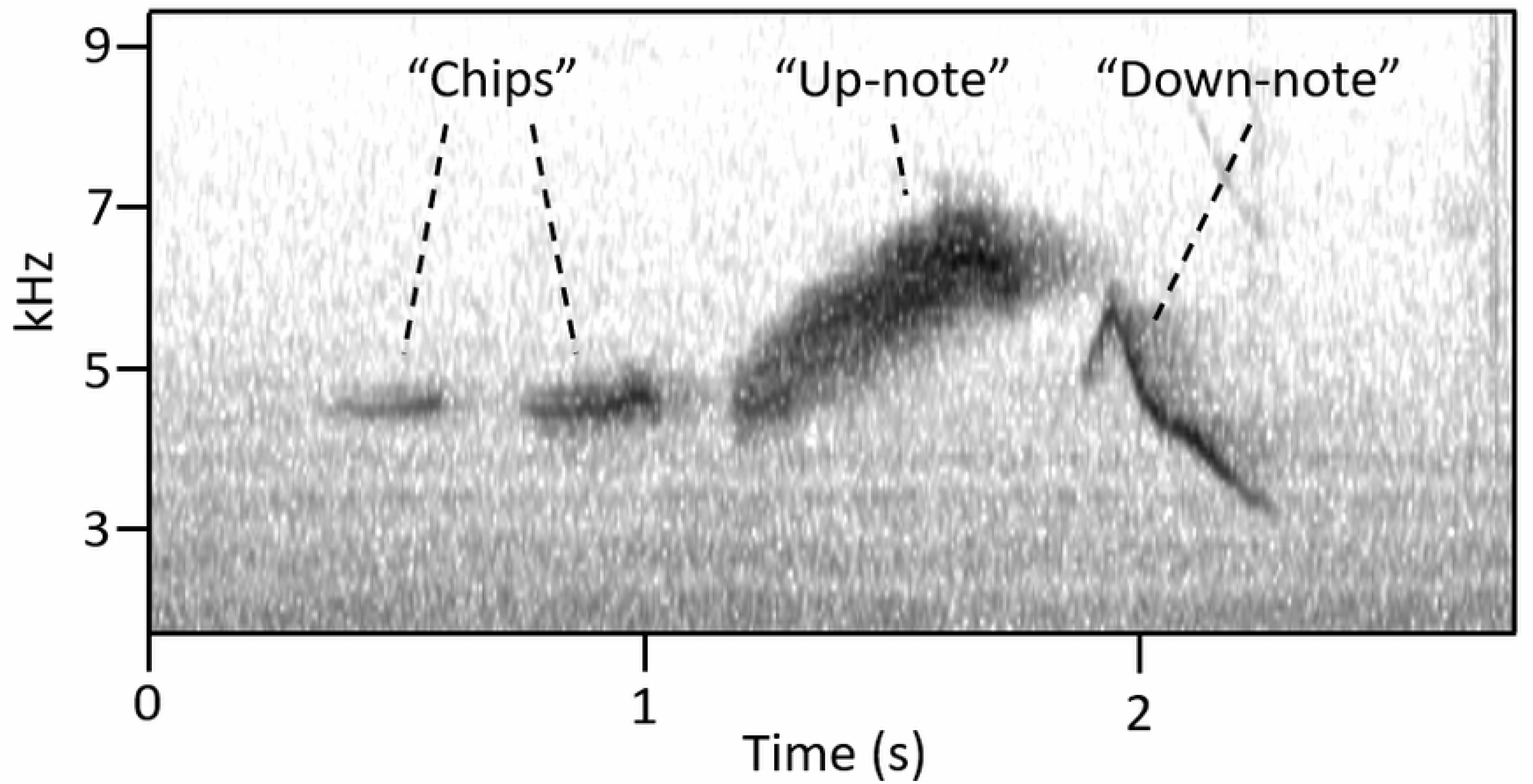
Spectrogram of a typical vitelline warbler song, with labeled chips, up-note, and down-note. All analyzed songs (n=417) contained one up-note. The up-note was preceded by 0-4 chips and followed by 0-1 down-notes. With these components, we recorded 10 basic song configurations. (Table 1, Fig 4). Down-notes were present in 249 songs and absent in 168 songs.

**Fig 4.**
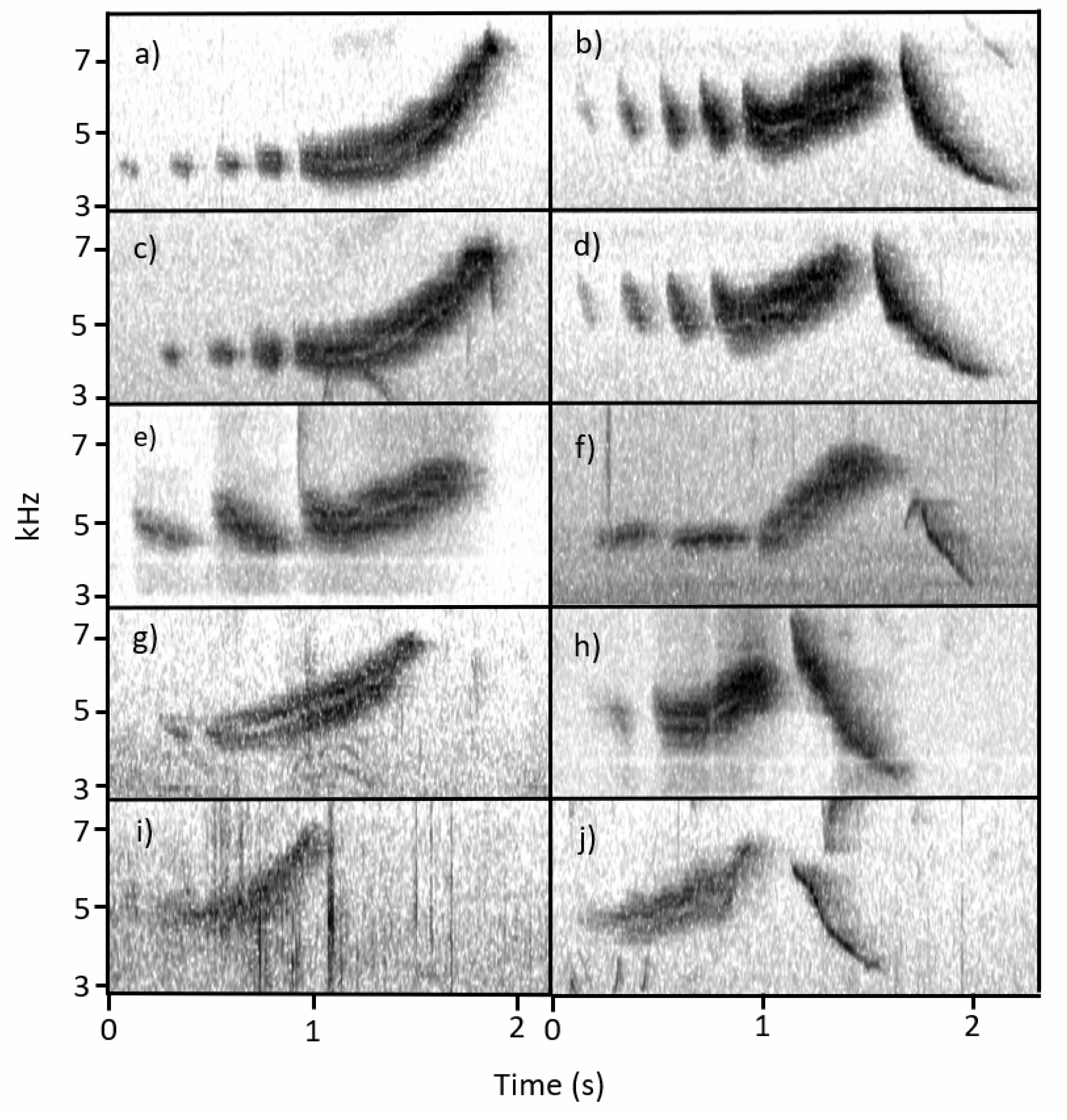
Ten representative song types of *S. vitellina*, categorized by number of components. Panels on the left (a, c, e, g, i) lack a down note and panels on the right (b, d, f, h, j) contain a down note. The top two panels (a and b) have four preceding chips, with each successive row below having one fewer chip.

**Table 1.** Components of 10 different song varieties, ordered by appearance in Fig 4.

| Chips (#) | Up-note (#) | Down-note (#) | Occurrence (n) | Figure 4 Identifier |
| --- | --- | --- | --- | --- |
| 4 | 1 | 0 | 20 | (a) |
| 4 | 1 | 1 | 9 | (b) |
| 3 | 1 | 0 | 54 | (c) |
| 3 | 1 | 1 | 91 | (d) |
| 2 | 1 | 0 | 61 | (e) |
| 2 | 1 | 1 | 74 | (f) |
| 1 | 1 | 0 | 22 | (g) |
| 1 | 1 | 1 | 57 | (h) |
| 0 | 1 | 0 | 11 | (i) |
| 0 | 1 | 1 | 18 | (j) |

Across all 10 song types, introductory chips had a mean frequency of 4.79 kHz, up-notes started at a mean frequency of 4.78 kHz and ended at 6.66 kHz, and down notes started at a mean frequency of approximately 6.62 kHz and ended at 3.78 kHz (Table 2).

**Table 2.** Frequency data for all components of 10 Vitelline Warbler song variations.

| Figure 4 identifier | Sample size | Chip frequency (kHz, mean ± SD) | | | | Up-note frequency (kHz, mean ± SD) | | Down-note frequency (kHz, mean ± SD) | |
| --- | --- | --- | --- | --- | --- | --- | --- | --- | --- |
|  |  | 4 | 3 | 2 | 1 | Start | End | Start | End |
| (b) | 9 | 4.99 ± 0.64 | 4.92 ± 0.51 | 4.90 ± 0.49 | 4.87 ± 0.44 | 5.08 ± 0.65 | 6.66 ± 0.39 | 7.23 ± 0.29 | 3.83 ± 0.32 |
| (a) | 20 | 4.64 ± 0.42 | 4.62 ± 0.40 | 4.64 ± 0.35 | 4.65 ± 0.35 | 4.67 ± 0.43 | 6.85 ± 0.38 | | |
| (d) | 91 | | 4.98 ± 0.43 | 4.97 ± 0.40 | 4.97 ± 0.38 | 5.05 ± 0.49 | 6.69 ± 0.38 | 6.79 ± 1.13 | 3.76 ± 0.35 |
| (c) | 54 | | 4.80 ± 0.37 | 4.84 ± 0.36 | 4.84 ± 0.36 | 4.87 ± 0.45 | 6.69 ± 0.56 | | |
| (f) | 76 | | | 4.64 ± 0.30 | 4.63 ± 0.31 | 4.70 ± 0.43 | 6.66 ± 0.46 | 6.56 ± 1.38 | 3.86 ± 0.50 |
| (e) | 59 | | | 4.65 ± 0.31 | 4.65 ± 0.33 | 4.72 ± 0.40 | 6.71 ± 0.52 | | |
| (h) | 57 | | | | 4.59 ± 0.46 | 4.60 ± 0.46 | 6.50 ± 0.54 | 6.29 ± 1.37 | 3.68 ± 0.46 |
| (g) | 22 | | | | 4.60 ± 0.46 | 4.60 ± 0.46 | 6.55 ± 0.44 | | |
| (j) | 18 | | | | | 4.62 ± 0.29 | 6.84 ± 0.57 | 6.83 ± 0.69 | 3.91 ± 0.79 |
| (i) | 11 | | | | | 4.49 ± 0.36 | 6.50 ± 0.48 | | |
| | Mean: | 4.75 ± 0.52 | 4.88 ± 0.43 | 4.78 ± 0.38 | 4.74 ± 0.39 | 4.78 ± 0.48 | 6.66 ± 0.48 | 6.63 ± 1.24 | 3.79 ± 0.47 |

Up-notes fell into four categories, all increasing in frequency from start to finish. The first shape variety, “check,” is characterized by a short initial decline in frequency followed by a longer rise in frequency (Fig 5a). The second variety, “convex,” is characterized by an initial steep rise in frequency, followed by a more gradual increase (Fig 5b). The third variety, “concave,” is characterized by a consistently low initial frequency that rises exponentially (Fig 5c). The fourth variety, “linear,” is characterized by a mostly consistent steepness as it increases steadily in frequency (Fig 5d).

**Fig 5.**
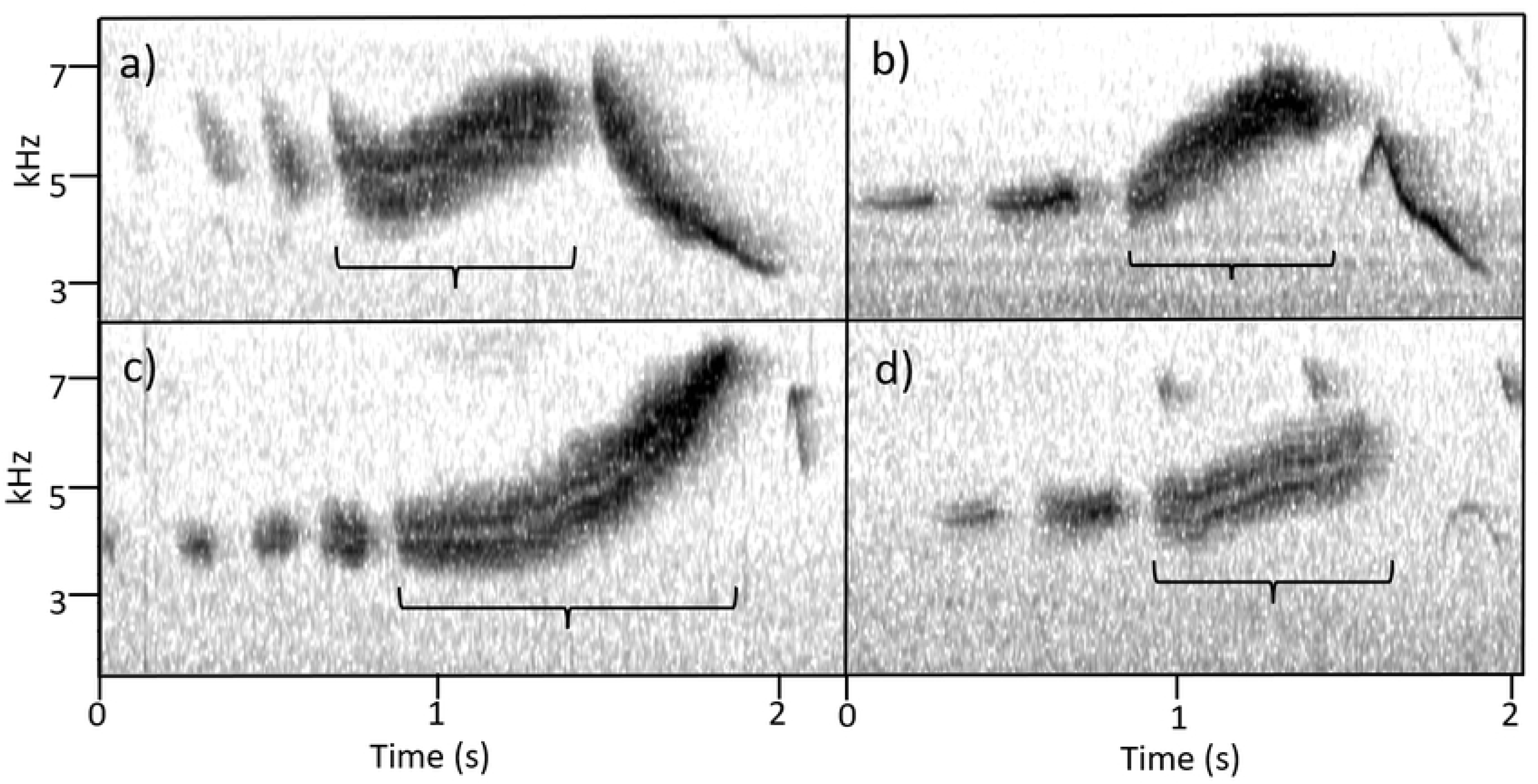
Four *S. vitellina* songs representing the array of up-note shapes. Shapes are labeled: (a) “check”, (b) “convex”, (c) “concave”, and (d) “linear”.

While up-note shapes were categorized qualitatively, quantitative analysis justified our categorization (Table 3). Among the four shapes, initial frequencies differed significantly (One-way ANOVA: F_3,416_ = 97.13, P<0.0001), as did end frequencies (One-way ANOVA: F_3,416_ = 34.98, P<0.0001). The starting frequency of “check” up-notes was higher than that of all three other shapes and the starting frequency of “linear” up-notes was higher than that of “convex” up-notes (Tukey’s Honestly Significant Difference Test, P<0.05 for each). The ending frequency of “concave” up-notes was higher than that of all three other shapes (Tukey’s Honestly Significant Difference Test, P<0.05 for each).

**Table 3.** Description of the frequency and length of the four up-note variations.

| | Frequency (kHz, mean ± SD) | | | Component length (s, mean ± SD) | | |
| --- | --- | --- | --- | --- | --- | --- |
| Shape | Start | Middle | End | Start-middle | Middle-end | Total |
| Check | 5.38 ±<br>0.31 | 4.97 ±<br>0.32 | 6.44 ± 0.43 | 0.12 ± 0.04 | 0.51 ±<br>0.10 | 0.64 ±<br>0.11 |
| Concave | 4.62 ±<br>0.37 | | 6.88 ±<br>0.38 | | | 0.79 ±<br>0.15 |
| Convex | 4.51 ±<br>0.47 | | 6.39 ±<br>0.54 | | | 0.74 ±<br>0.16 |
| **Linear** | 4.71 ±<br>0.34 | | 6.51 ±<br>0.49 | | | 0.68 ±<br>0.17 |

Up-notes of the “check” shape decrease in frequency before rising, so frequency data is represented for the start, low-point (“middle”) and end of each note.

Down notes fell into five general shape categories, with all ending at a lower frequency than they started (Fig 6). The first variety, “concave,” was characterized by an initial steep drop in frequency that attenuates to a near constant frequency. The second variety, “linear,” was qualitatively similar to concave, but was characterized by a more consistent decline in frequency. The third variety, “parabolic,” was characterized by an initial short rise in frequency, followed by a longer drop in frequency. The fourth variety, “hyperbolic X,” was characterized by relatively shallow declines in frequency at the beginning and end, with a steep decline in the middle of the note. The fifth variety, “hyperbolic Y,” was characterized by steep declines at the start and end of the note, with a shallower decline in the middle.

**Fig 6.**
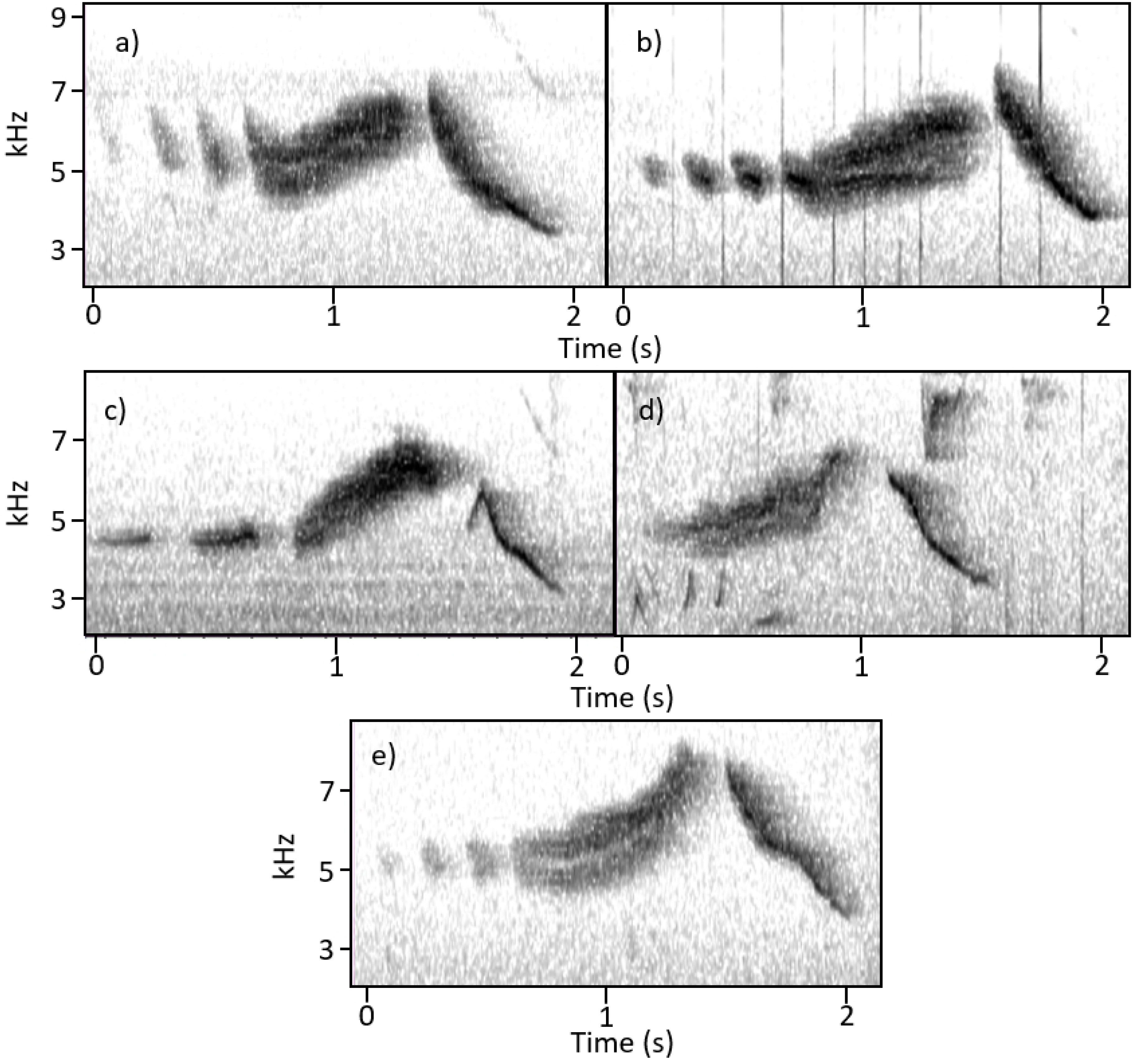
Five Vitelline Warbler songs representing the array of down-note shapes. Shapes are labeled “concave” (a), “linear” (b), “parabolic” (c), “hyperbolic X” (d), and “hyperbolic Y” (e).

In addition to their qualitative differences, down-note shapes differed quantitatively (Table 4). We found significant differences in the starting frequencies (One-way ANOVA: F_4,249_ = 148.15, p<0.0001) and end frequencies (One-way ANOVA: F_4,249_ = 19.96, p<0.0001) of the five shapes. The starting frequency of “parabolic” down-notes was lower than that of all four other shapes and the starting frequency of “hyperbolic X” down-notes was lower than that of the remaining three shapes (Tukey’s Honestly Significant Difference Test, p<0.05 for each). The ending frequency of “parabolic” down-notes was lower than that of all shapes except “hyperbolic X,” and the ending frequency of “concave” down-notes was lower than that of both “hyperbolic Y” and “linear” down-notes (Tukey’s Honestly Significant Difference Test, p<0.05 for each).

**Table 4.** Description of the frequency and length of five down-note variations.

| Shape | Frequency (kHz, mean ± SD) | | | Component length (s, mean ± SD) | | |
| --- | --- | --- | --- | --- | --- | --- |
|  | Start | Middle | End | Start-middle | Middle-end | Total |
| **Concave** | 7.05 ± 0.70 | | 3.70 ± 0.29 | | | 0.49 ± 0.10 |
| **Hyperbolic X** | 5.88 ± 0.04 | | 3.68 ± 0.22 | | | 0.33 ± 0.06 |
| **Hyperbolic Y** | 7.10 ± 0.61 | | 4.10 ± 0.37 | | | 0.42 ± 0.09 |
| **Linear** | 7.12 ± 0.74 | | 3.99 ± 0.55 | | | 0.41 ± 0.09 |
| **Parabolic** | 4.19 ± 0.54 | 4.83 ± 0.47 | 3.38 ± 0.45 | 0.06 ± 0.02 | 0.38 ± 0.17 | 0.44 ± 0.18 |

Down-notes of the "parabolic" shape rise in frequency before declining, so frequency data is represented for the start, high-point ("middle") and end of each note.

### Call characteristics of the Vitelline Warbler

There appears to be no published description of the calls of *S. vitellina* (12). Here, we provide descriptions of calls we observed and recorded.

During one survey, we observed a male *S. vitellina* calling at close range. We recorded the warbler’s vocalizations for 88 seconds, over the course of which it produced 26 short call notes. All 26 call notes qualitatively appeared to be the same vocalization. These vocalizations lasted between 0.029 and 0.047 seconds (mean ± SD = 0.037 ± 0.004s). Vocalizations started at a higher frequency (mean ± SD = 6.892 ± 0.094 kHz) and ended at a lower frequency (mean ± SD = 5.811 ± 0.114 kHz). When this same male sang, the songs’ chips were at a lower frequency (mean ± SD = 4.97 ± 0.12 kHz), suggesting that calls (not part of a song) and chips (part of a song) are different vocalizations.

## Discussion

Our findings demonstrate variation in both song composition and individual song elements within this population of *S. vitellina*. Specifically, existence of the down-note and the variety in shapes of up-notes and down-notes represents a level of complexity with no analog in the repertoire of *S. discolor*, by far the closest relative of *S. vitellina*.

There are multiple possible mechanisms for the increased complexity in song of the island-restricted *S. vitellina* as compared to its migratory *S. discolor* relative. High levels of song variation have been observed in other island-restricted songbirds, with proposed mechanisms including reduced selection for song specificity (17) and the non-adaptive accumulation of song errors (deviations from typical songs) that do not greatly influence individual fitness (18). Furthermore, island bird populations can experience “character release,” whereby relatively few competing species allows a trait (in this case song) to become more variable, due to less selective pressure for specialization (19). Apparently, it is not inevitable or even common for there to be divergence in the song repertoire between island and mainland populations of songbirds. Morinay et al. (20) compared 49 pairs of island and mainland passerine species and found no general pattern of higher or lower complexity in island species.

It remains unclear whether differences between *S. discolor* and *S. vitellina* are due to evolution of singing behavior in the island species, the migratory species, or both. The last common ancester of *S. discolor* and *S. vitellina* has been estimated at 1.2 million years (11). The current song of *S. discolor* is relatively simple (21), indicating that in the 1.2 million years since their split, either the migratory *S. discolor* song has simplified, or the island-resident *S. vitellina* song has become more complex. Further study of these two warblers, particularly the understudied *S. vitellina,* could help shed light on the mechanisms by which bird song evolves.

### Conservation Implications

More studies of *S. vitellina* are needed, given a variety of environmental threats facing the Cayman and Swan Islands. At least three significant threats face *S. vitellina* today, including feral cats (22,23), climate change, and increasing development on the island (16). Conservation efforts are complicated by *S. vitellina’s* relative obscurity. Despite the fact that the vast majority of *S. vitellina* are found in the Cayman Islands, they receive little national attention, likely due to their perceived ubiquity and lack of visually striking features compared to the more charismatic Cuban Parrot (*Amazona leucucephala*) (24,25), Red-footed Booby (*Sula sula*) (26), and Brown Booby (*Sula leucogaster*) (12,27).

The poorly-documented life history of *S. vitellina* warrants further research and monitoring. There is a need for better knowledge of habitat and food requirements, the extent and severity of feral cat predation, reproductive success and demography, and population densities across the species’ restricted range. A more complete understanding of this endemic songbird’s behavior and ecology would have value for targeted conservation strategies. Furthermore, small islands like Little Cayman offer unique opportunities to address general questions regarding evolution of birdsong.

## Acknowledgements

We thank Victoria Moss for her assistance in conducting distribution surveys and analyzing recordings. We would also like to thank Matthew Ayres, Chris Rimmer, and Miranda Zammarelli for their thorough revisions of draft manuscripts. We thank Pooja Panwar for offering advice on the presentation of results. Finally, we would like to thank the staff of the Central Caribbean Marine Institute and all other residents of Little Cayman for welcoming and hosting us for the duration of this project.

